# Analysis of expression Quantitative Trait Loci for *NLGN4X* in relation to language and neurodevelopmental function: an exploratory analysis using FUSION and GTEx workflows

**DOI:** 10.1101/2021.06.29.450314

**Authors:** Nuala H. Simpson, Dorothy V.M. Bishop, Dianne F. Newbury

## Abstract

Language disorders in children are highly heritable, but progress in identifying genetic variants that contribute to language phenotypes has been slow. Here we applied a novel approach by identifying SNPs that are associated with gene expression in the brain, taking as our focus a gene on the X chromosome, *NLGN4X*, which has been postulated to play a role in neurodevelopmental disorders affecting language and communication. We found no significant associations between expression quantitative trait loci (eQTLs) and phenotypes of nonword repetition, general language ability or neurodevelopmental disorder in two samples of twin children, who had been selected for a relatively high rate of language problems. We report here our experiences with two methods, FUSION and GTEx, for eQTL analysis. It is likely that our null result represents a true negative, but for the interest of others interested in using these methods, we note specific challenges encountered in applying this approach to our data: a) complications associated with studying a gene on the X chromosome; b) lack of agreement between expression estimates from FUSION and GTEx; c) software compatibility issues with different versions of the R programming language.

## Introduction

Expression quantitative trait loci (eQTLs) are single nucleotide polymorphisms (SNPs) that are associated with variations in the expression level of a gene i.e. variations in the genetic sequence that are correlated with the amount of messenger RNA a given gene produces (Nica and Dermitzakis, 2013). This correlation can reflect a functional role for the given variant (e.g. by transcription factor binding, DNA methylation and histone modifications) or, alternatively, may indicate that the given variant is in linkage disequilibrium (LD) with a functional variant (Pai et al., 2015). These loci therefore represent good candidates for the subtle alteration of gene regulation which may be important in modifier effects and complex disorders. However, the identification and mapping of eQTLs requires information regarding genetic sequence variations linked to gene expression across tissues and over time and is therefore complex to investigate at an individual level.

In this paper, we evaluate the use of existing datasets and computational algorithms to investigate expression levels of a *neuroligin* gene, *NLGN4X*, within the brain in correlation with three language phenotypes in a sample of twin children, recruited to over-select cases with language problems.

### The *neuroligin* hypothesis

Previously we had studied *neuroligins* in a sample of children with an extra X or Y chromosome (sex chromosome trisomies) (Newbury et al, 2020). We had hypothesised a ‘double hit’ model, whereby elevated levels of *NLGN4X/Y* in those with an extra sex chromosome might play a role in the modification of language and neurodevelopmental phenotypes of sex chromosome trisomies (Bishop and Scerif, 2011). We did not find support for our predictions, but we did not test *NLGN4X/Y* expression directly: rather we looked for increased associations in this sample between language/neurodevelopmental phenotypes and autosomal variants in genes that interact with *neuroligins*.

The notion that *neuroligins* may be implicated in the neurodevelopmental problems of children with sex chromosome trisomies was, however, supported by research from another group who directly measured *NLGN4Y* gene expression in blood, and showed a correlation between the expression of this gene and social responsiveness and autism symptoms in a small sample of 47,XYY males (Ross et al., 2015).

The development of computational algorithms to measure gene expression makes it feasible to test the *neuroligin* hypothesis by inferring individual differences in expression of *NLGN4X* using sequence variants and looking for association with language and neurodevelopmental phenotypes. The sample of children with sex chromosome trisomies is not suitable for this analysis because the reliable calling and imputation of SNPs on the X chromosome in sex chromosome aneuploidy cases is difficult. However, the computational algorithms can be applied to a comparison sample of twin children, enriched for cases of language difficulties, who were a comparison group in our studies of sex chromosome trisomies, and had been assessed on the same phenotypic measures.

Analysis of eQTLs is still a relatively new method and there is no agreement on the optimal approach. Computational methods have mostly been designed for large-scale transcriptome-wide analysis studies (TWAS). We were interested in exploring the potential of applying this approach to a candidate gene, using a relatively small sample. We applied and compared two alternative workflows, FUSION (Gusev et al., 2016) and GTEx (https://gtexportal.org/home/), to identify eQTL-panels for *NLGN4X* expression, both of which use publicly available expression and genotype datasets to identify correlations between specific genetic sequence variants and *NLGN4X* expression in the brain. The derived panels were then used to infer gene expression levels in our independent sample for whom comprehensive language/neurodevelopmental phenotypic measures and genotype data were available. This allowed us to evaluate the hypothesis that *NLGN4X* expression levels are correlated with language and neurodevelopmental phenotypes.

## Methods

### Twin datasets

The twin dataset has been described in detail previously (Wilson and Bishop, 2018). In summary, the children were aged from 6 years 0 months to 11 years 11 months, were taking part in a twin study of language and laterality, had completed the same test battery, and had English as a first language at home. The aim had been to over-recruit twin pairs in which one or both twins had language or literacy problems. This was coded on the basis of parental response on a telephone interview: any mention of language delay, history of speech and language therapy, current language problems or dyslexia was coded as ‘parental concern’. For the current analysis, we grouped together all twins, regardless of zygosity and parental concern, and then divided them into two subsamples by selecting one twin from each pair at random, after excluding 18 cases with missing or insufficient DNA. Twin set 1 contained 184 individuals (91 males and 93 females) and Twin set 2 contained 186 individuals (100 males and 86 females). This meant we could replicate the analysis for twins with a diploid (typical) karyotype. Note that this replication sample is not independent, as the genotype for the MZ twins is the same in the two subsamples, and is related for DZ twins.

### Phenotypic measures

We analysed three quantitative phenotypes that have been described in more detail previously (Newbury et al., 2020): nonword repetition, language factor and global neurodevelopment score. They represent increasingly general indices of language/communication problems. Nonword repetition, was used as a measure of a specific component of language processing, phonological short-term memory, which is a heritable marker of Developmental Language Disorder (Bishop et al., 1996). A general language factor was derived from the language tests: Verbal Comprehension, Oromotor Sequences, Sentence Repetition and Vocabulary. The global neurodevelopment score was created using all of the available information from a parental report and extends beyond language to include features of other neurodevelopmental disorders, including intellectual disability, attentional problems and autistic features. For all three phenotypes a low score indicates impairment.

### Workflows

Two workflows were applied to the identification of eQTLs within publically available gene expression datasets (see Figure 1). The first employed the FUSION algorithm (see FUSION workflow below) and the second directly extracted information from the GTEx (Lonsdale et al., 2013) web portal (GTEx workflow). Each identified eQTL-panel was then used to infer *NLGN4X* expression levels within two twin datasets, allowing comparison and evaluation of inferred expression levels across samples. FUSION-inferred expression levels were then used to investigate correlations between *NLGN4X* expression and language ability.

**Figure 1.**
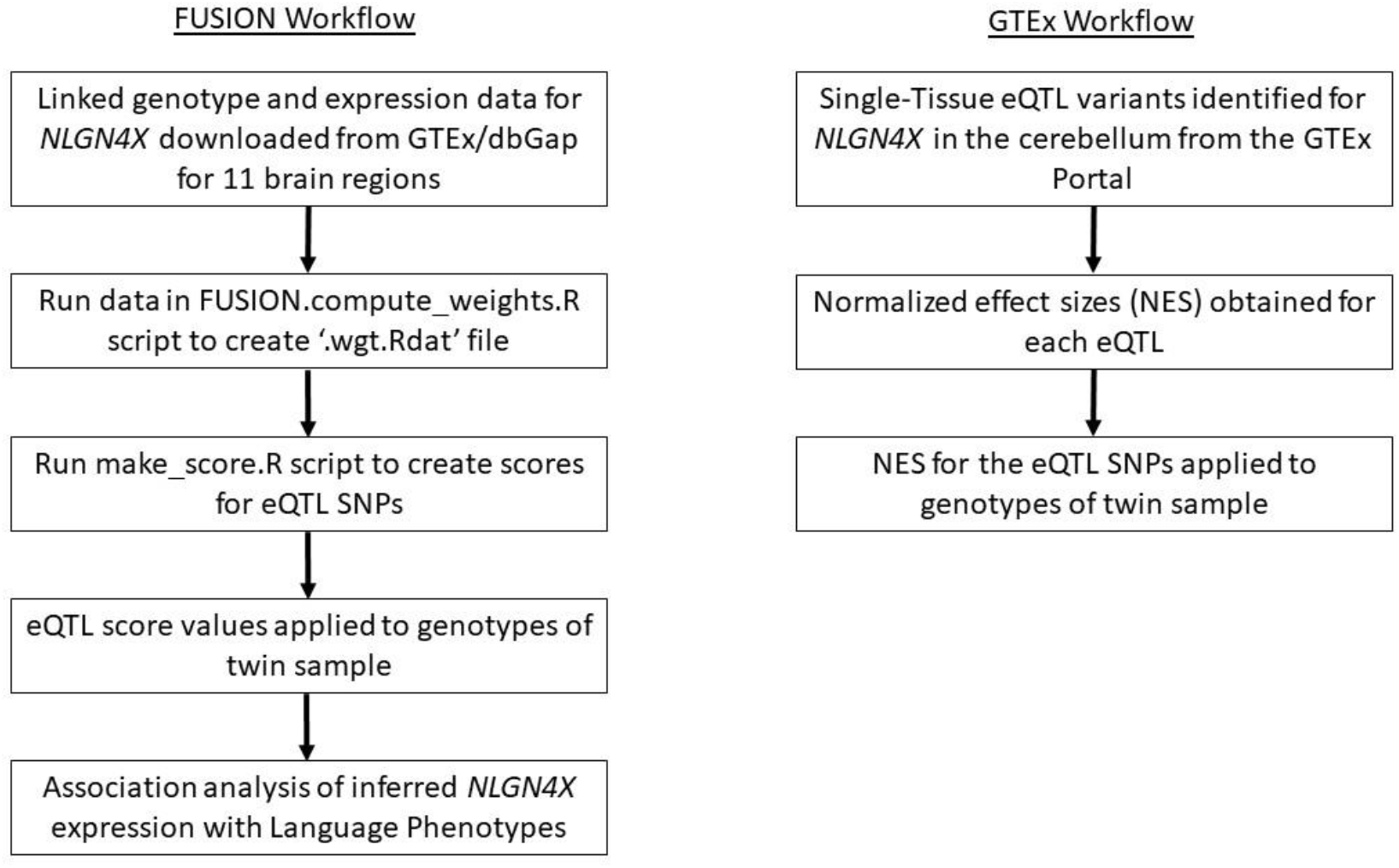
Workflows for the FUSION and GTEx methods.

### FUSION workflow - Extraction of genotype and expression data

Gene expression levels and genotypes were first obtained from an independent database where information on expression of genes in different tissues is available. For this step, linked genotype and gene expression data were downloaded from GTEx/dbGaP (Genotype-Tissue Expression (GTEx (https://gtexportal.org/home/)) (phs000424.v8.p2) following access and data transfer agreements. Genotype files were obtained for 450 individuals (158 females and 292 males) for *NLGN4X* (gene position chrX: 5808083-6146706 (hg19)) plus 500kb 5’ and 3’ to this gene. Expression data for a subset of these individuals was available across 11 brain tissues (Amygdala, anterior cingulate cortex, caudate basal ganglia, cerebellar hemisphere, cerebellum, cortex, frontal cortex (BA9), hippocampus, hypothalamus, nucleus acumbens basal ganglia and putamen basal ganglia). Expression data was available for between 18 and 72 individuals per brain tissue. Genotypes were pruned to contain only SNPs present in the 1000 genomes LD reference data and were forced in Plink (Purcell et al., 2007) to have the same reference allele as the 1000 genomes. This was a requirement for the FUSION script to run.

### Identification of eQTLs using FUSION

FUSION employs a two-step method within the R-platform v3.4.1 (R Core Team, 2013). Genotype and associated expression data from GTEx/dbGaP were linked through the FUSION.compute_weights.R script which identified potential eQTL variants. If an eQTL is identified a ‘.wgt.RDat’ file is created. A second Rscript (make_score.R) then uses this file to weight the eQTLs producing a score-file (weighted scores per SNP), which can then be applied to other datasets with genotype data for overlapping or related SNPs to infer gene expression levels (Gusev et al., 2016) (http://gusevlab.org/projects/fusion/). The scores per SNP were used to infer expression levels in the twin datasets. The FUSION R scripts were run separately for the female and male GTEx genotype files for each of the 11 brain regions studied.

The accurate generation of a weighting file relies upon an adequate sample size and a consistent relationship between SNP genotype and expression levels of the gene in question. Of the 11 brain regions studied in the FUSION workflow, FUSION was able to identify *NLGN4X* eQTLs for only one brain tissue in females (cerebellum) and one region in male samples (hippocampus).

### GTEx workflow - Extraction of *NLGN4X* eQTLs from the GTEx database

The second inference workflow was manually curated and used information directly from the GTEx Portal (https://gtexportal.org/home/) to consider the correlation between genotype and *NLGN4X* expression. Normalized effect sizes (NES) were manually obtained from the GTEx Portal for the brain tissue cerebellum. The cerebellum was the tissue identified by the FUSION workflow to contain eQTLs for *NLGN4X* in females and was relevant for our purposes, given its role in language function (Marien et al., 2014). Therefore eQTLs in this brain tissue were analysed in the GTEx workflow as a comparison to the FUSION eQTL-panel.

The NES values represent the slope of the linear regression line between gene expression associated with the reference allele (i.e. the form of the genetic variant found in the Human Reference Genome) and alternative alleles of a given eQTL SNP variant. The magnitude of these values has no direct biological interpretation but differences between alternative and reference alleles represent a correlation that may indicate the presence of an eQTL. All significant Single-Tissue eQTL variants (Data Source: GTEx Analysis Release V8 (dbGaP Accession phs000424.v8.p2) downloaded on 01/10/2019) listed in the GTEx portal for *NLGN4X* in the cerebellum were identified and the NES used as a weighting value for gene expression changes associated with the presence of the alternative allele. The NES values will not explain all gene expression but are used as an indicator of expression level.

### Inference of gene expression levels in the twin dataset

Genotype data for all variants within each eQTL panel was extracted from genomewide SNP data that had been imputed for the twin dataset using the Michigan Imputation server (Das et al., 2016) as described previously (Newbury et al., 2020). Genotype files for the twin dataset were split into males (191 individuals) and females (179 individuals). eQTL SNPs that had not been genotyped or imputed in the twins, were replaced by proxies identified using linkage disequilibrium information derived from CEU (UTAH Residents from North and West Europe) and GBR (British in England and Scotland) populations in the website https://ldlink.nci.nih.gov. Proxy SNPs were selected to be physically close and in high LD (r^2^>0.8) to the original variant.

Prior to inference of gene expression, each eQTL panel was subject to a number of quality control checks performed in VCFtools (Danecek et al., 2011) and PLINK (Purcell et al., 2007). Genotype quality was assessed by the examination of imputation R-squared values. There were no parental genotypes available for the twins and so Mendelian inheritance could not be checked. All genotyping rates were 100% for genotyped and imputed eQTL SNPs. Minor allele frequencies (--freq), Hardy Weinberg Equilibrium probabilities (--hardy) and pairwise linkage disequilibrium (--r2) were calculated within PLINK (Purcell et al. 2007) for all eQTLs.

For each eQTL SNP the weighted score value from the FUSION workflow or the NES value from the GTEx workflow associated with the minor alleles were summed to create a cumulative total across all eQTL SNPs in each twin individual and so infer expression levels.

## Results

### FUSION workflow - Adaptation for analysis of X chromosome gene

The FUSION software does not allow the analysis of X chromosome annotated SNPs, as these may differ in dosage between males and females. For genes that are X-inactivated dosage should be similar for the two sexes, but some genes escape X-inactivation such as *NLGN4X*. In this study, sex-specific analyses had to be forced by assigning *NLGN4X* SNPs to an autosome in the input files. Since all females have two copies of the X chromosome, this SNP reassignment simply allowed the software to recognise the data. For males, this reassignment made the (incorrect) assumption that two alleles will be present for all SNPs. The software treated males as always being homozygous and so estimates of gene expression levels assigned to an allele will be inexact. However, since this estimate can be assumed to be the same for all males, the relationships between gene expression levels will remain consistent.

Of the 726 SNPs examined in the FUSION workflow, fifteen were identified as having an effect on *NLGN4X* gene expression in the cerebellum in females (Table 1 and Figure 2). Eleven of these eQTLs fell 3’ to the *NLGN4X* transcript across positions chrX: 5462650-5788968 (hg19) (Table 1 and Figure 2) coinciding with mapped histone marks and extending across a neighbouring transcript *RP11-733018*. Two eQTLs fell 5’ of *NLGN4X* across positions chrX: 6,226,090-6,446,854 (hg19) (Table 1 and Figure 2) flanking a neighbouring miRNA (*miR4770*). Two eQTLs fell within the *NLGN4X* gene itself, both towards the 3’ end of the gene (chrX: 5,816,328-5,825,728, hg19) (Table 1) around exon 5 of 6 (referring to NM_181322).

**Figure 2:**
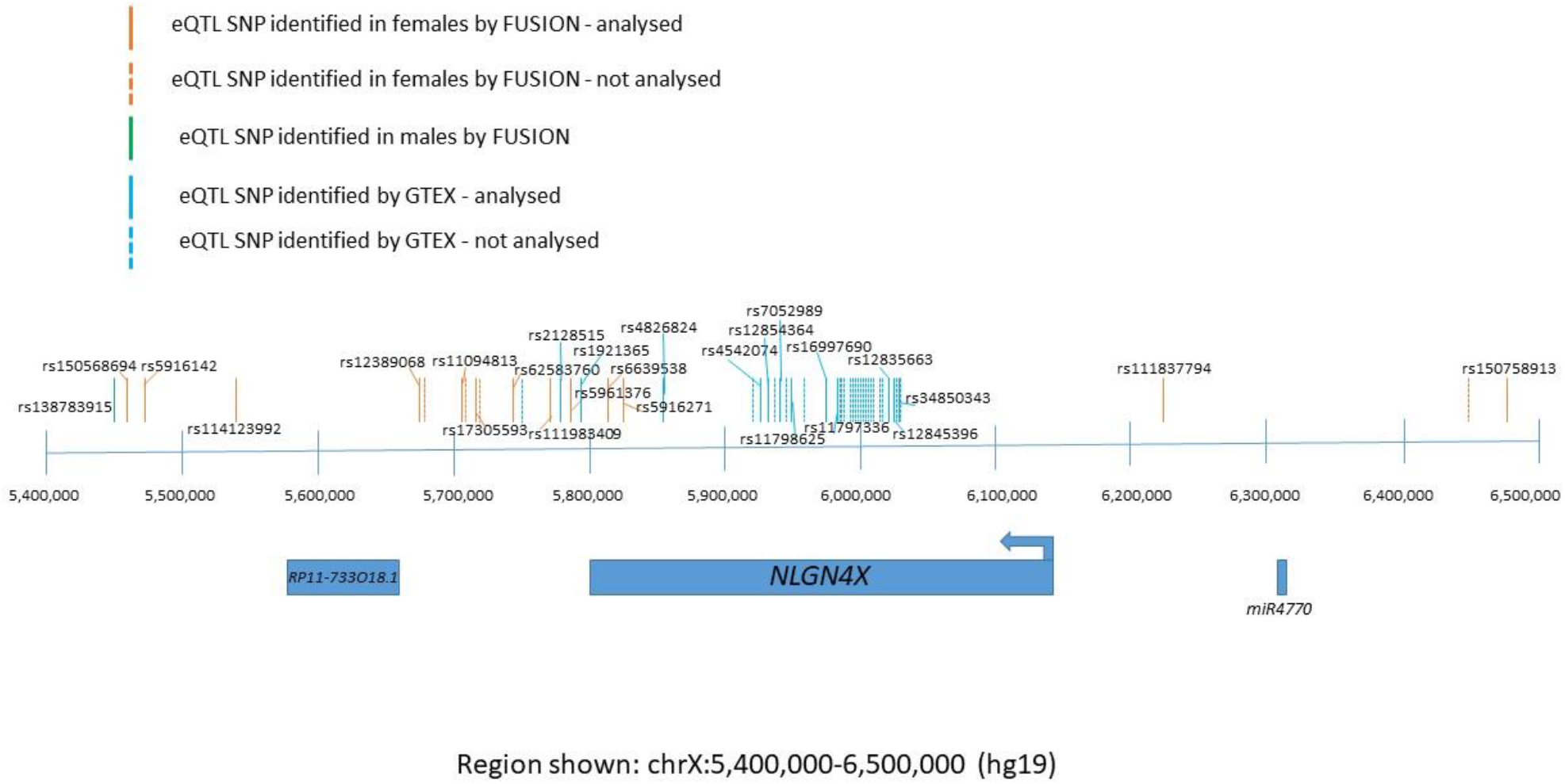
Schematic of the eQTL SNPs identified.

**Table 1.** *NLGN4X* eQTL-SNP panel Identified by FUSION workflow in females in the cerebellum.

| SNP name | position (hg19) | Minor allele | Major allele | Expression change associated with minor allele | Present in female twin genotypes? | Imputed or genotyped? | R-squared value from info for imputed SNPs (above 0.7 acceptable) | Allele frequencies from UCSC | SNP used in final analysis? |
| --- | --- | --- | --- | --- | --- | --- | --- | --- | --- |
| rs150568694 | X:5462650 | T | C | -0.000729 | Yes | Imputed | 0.98 | T: 2.755% ;<br>C: 97.245% | Yes |
| rs5916142 | X:5474819 | A | G | -0.067126 | Yes | Imputed | 0.86 | A: 7.974% ;<br>G: 92.027% | Yes |
| rs114123992 | X:5543547 | A | G | 0.025407 | Yes | Imputed | 0.81 | A: 9.192% ;<br>G: 90.808% | Yes |
| rs12389068 | X:5671225 | T | C | -0.006136 | Yes | Imputed | 0.95 | T: 45.642% ;<br>C: 54.358% | Yes |
| rs5961359 | X:5679841 | A | C | -0.000186 | Yes | Imputed | 0.96 | A: 44.689% ;<br>C: 55.311% | SNP removed due to high LD |
| rs12556975 | X:5704309 | T | C | -0.165558 | No | No | na | T: 8.291% ;<br>C: 91.709% | No |
| rs17305593 | X:5713122 | A | G | -0.005017 | Yes | Imputed | 0.98 | A: 8.265% ;<br>G: 91.735% | Yes |
| rs73182556 | X:5723303 | T | C | -0.000152 | Yes | Imputed | 0.97 | T: 8.238% ;<br>C: 91.762% | SNP removed due to high LD |
| rs62583760 | X:5743758 | A | G | 0.075786 | Yes | Imputed | 0.83 | A: 10.066% ;<br>G: 89.934% | Yes |
| rs111983409 | X:5773454 | A | G | -0.087599 | Yes | Imputed | 0.96 | A: 16.080% ;<br>G: 83.921% | Yes |
| rs5961376 | X:5788968 | T | C | -0.051464 | Yes | Imputed | 0.99 | T: 18.146% ;<br>C: 81.854% | Yes |
| rs6639538 | X:5816328 | G | A | 0.020656 | Yes | Imputed | 0.99 | G: 36.636% ;<br>A: 63.364% | Yes |
| rs5916271 | X:5825728 | C | A | 0.144361 | Yes | Imputed | 0.96 | C: 12.795% ;<br>A: 87.205% | Yes |
| rs111837794 | X:6226090 | G | T | 0.016388 | Yes | Imputed | 0.99 | G: 21.907% ;<br>T: 78.093% | Yes |
| rs182391645 | X:6446854 | T | C | -0.301837 | No | No | na | T: 2.252% ;<br>C: 97.748% | No |
| rs11094813 | X:5704257 | A | G | -0.165558 | Yes | Imputed | 0.99 | A: 8.238% ;<br>G: 91.762% | Yes - proxy for rs12556975 |
| rs150758913 | X:6484590 | A | T | -0.301837 | Yes | Imputed | 0.54 | A: 2.649% ;<br>T: 97.351% | Yes - proxy for rs182391645 |

In male samples, one of the 726 SNPs examined in the FUSION workflow was identified to be correlated with *NLGN4X* expression in the hippocampus (rs138783915) (Table 2 and Figure 2). This SNP fell 355kb 3’ (chrX: 5452531, hg19) of the *NLGN4X* gene and did not overlap with any of the eQTLs identified in females.

**Table 2.** *NLGN4X* eQTL-SNP panel Identified by FUSION workflow in males in the hippocampus.

| SNP name | position (hg19) | Minor allele | Major allele | Expression change associated with minor allele | Present in male twin genotypes? | Imputed or genotyped? | R-squared value from info for imputed SNPs (above 0.7 acceptable) | Allele frequencies from UCSC |
| --- | --- | --- | --- | --- | --- | --- | --- | --- |
| rs138783915 | X:5452531 | T | C | 3.26 | yes | Imputed | 0.72 | T: 0.556% ;<br>C: 99.444% |

The SNP rs138783915 had been imputed in the twin dataset but during quality checks, it was noted that this variant had a minor allele frequency of 0.5%. Inspection of the genotype data showed that only two twin male individuals carried the minor allele for this variant. As the eQTL identified in male hippocampal samples relied upon this single SNP alone (Table 2), the available twin sample size was deemed too small to allow the accurate inference of gene expression and this variant and brain tissue were therefore removed from further analyses meaning that *NLGN4X* expression was not inferred in the male twin datasets.

Figure 2 shows all of the eQTL SNPs identified by the FUSION and GTEx workflows.

### GTEx workflow - Inference Process

The inference of gene expression using FUSION was a new technique to us. In addition, there were many complications regarding the use of X chromosome data in this script. As an independent validation approach, we therefore used information directly from the GTEx Portal (https://gtexportal.org/home/) to consider the correlation between genotype and *NLGN4X* expression. This method was not as sophisticated as FUSION and did not generate a weighted model of gene expression. Rather, we considered the reported correlation between each variant and gene expression to generate an additive eQTL set.

Thirty four eQTL SNPs were identified in the GTEx database that correlated with *NLGN4X* expression in the cerebellum (Table 3 and Figure 2). These eQTLs were not specified to be gender specific, unlike the FUSION output, which was analysed separately for males and females.

**Table 3.** *NLGN4X* eQTL-SNP panel identified by the GTEx workflow.

| SNP name | position (hg19) | Allele frequencies from UCSC | Imputed or genotyped | Normalized effect size | P-Value | SNP used in final analysis? |
| --- | --- | --- | --- | --- | --- | --- |
| rs5916251 | X:5753795 | C: 91.258% ; T: 8.742% | Imputed | -0.21 | 0.00011 | No |
| **rs2128515** | **X:5777815** | **A: 8.715% ; G: 91.285%** | **Imputed** | **-0.21** | **0.00011** | **Yes** |
| **rs1921365** | **X:5789410** | **A: 28.450% ; G: 71.550%** | **Imputed** | **-0.16** | **0.00008** | **Yes** |
| **rs4826824** | **X:5859733** | **C: 11.550% ; G: 88.450%** | **Imputed** | **-0.21** | **0.0000079** | **Yes** |
| rs35812438 | X:5927129 | C: 85.616% ; T: 14.384% | Imputed | 0.18 | 0.00011 | No |
| **rs4542074** | **X:5930024** | **A: 8.556% ; G: 91.444%** | **Genotyped** | **0.22** | **0.000038** | **Yes** |
| **rs12854364** | **X:5933655** | **C: 90.172% ; T: 9.828%** | **Imputed** | **0.19** | **0.000059** | **Yes** |
| rs12855992 | X:5933656 | A: 89.775% ; G: 10.225% | Imputed | 0.19 | 0.000059 | No |
| **rs7052989** | **X:5944178** | **A: 22.252% ; G: 77.748%** | **Imputed** | **0.18** | **0.000061** | **Yes** |
| rs6639583 | X:5949912 | C: 64.238% ; T: 35.762% | Imputed | -0.16 | 0.00004 | No |
| **rs11798625** | **X:5950034** | **G: 80.636% ; T: 19.364%** | **Imputed** | **0.17** | **0.000091** | **Yes** |
| rs35620385 | X:5959074 | C: 81.828% ; T: 18.172% | Imputed | 0.19 | 0.000014 | No |
| **rs16997690** | **X:5977582** | **C: 82.305% ; T: 17.695%** | **Imputed** | **0.2** | **0.000007** | **Yes** |
| **rs11797336** | **X:5980065** | **G: 18.305% ; T: 81.695%** | **Imputed** | **0.2** | **0.000007** | **Yes** |
| rs12840457 | X:5981472 | A: 12.927% ; G: 87.073% | Imputed | 0.22 | 0.0000014 | No |
| rs12848943 | X:5981538 | G: 18.146% ; T: 81.854% | Imputed | 0.19 | 0.000014 | No |
| rs34678738 | X:5982482 | A: 17.775% ; G: 82.225% | Imputed | 0.19 | 0.000014 | No |
| rs12851920 | X:5991222 | C: 18.464% ; T: 81.536% | Imputed | 0.18 | 0.000029 | No |
| rs7881769 | X:5992822 | C: 17.801% ; T: 82.199% | Imputed | 0.19 | 0.000012 | No |
| rs11797625 | X:5992847 | C: 13.139% ; T: 86.861% | Imputed | 0.21 | 0.0000034 | No |
| rs12836548 | X:5999052 | C: 17.589% ; G: 82.411% | Imputed | 0.19 | 0.000024 | No |
| rs10522048 | X:5999991 | C: 18.199% ; G: 81.801% | Imputed | 0.19 | 0.000013 | No |
| rs11795639 | X:6003984 | C: 82.437% ; T: 17.563% | Imputed | 0.19 | 0.000024 | No |
| rs11798311 | X:6004432 | A: 17.616% ; T: 82.384% | Imputed | 0.19 | 0.000024 | No |
| rs7060929 | X:6008597 | A: 82.384% ; G: 17.616% | Imputed | 0.19 | 0.000024 | No |
| rs73627050 | X:6012628 | C: 82.675% ; G: 17.324% | Imputed | 0.19 | 0.000024 | No |
| rs11796499 | X:6012960 | C: 17.510% ; G: 82.490% | Imputed | 0.17 | 0.00012 | No |
| rs9698643 | X:6019890 | A: 60.000% ; G: 40.000% | Imputed | 0.19 | 0.000024 | No |
| rs12008603 | X:6021181 | A: 82.464% ; C: 17.536% | Imputed | 0.19 | 0.000012 | No |
| **rs12835663** | **X:6027469** | **C: 14.172% ; T: 85.828%** | **Imputed** | **0.2** | **0.0000075** | **Yes** |
| **rs12845396** | **X:6029533** | **A: 67.788% ; T: 32.212%** | **Imputed** | **0.19** | **0.0000021** | **Yes** |
| rs12839271 | X:6031466 | C: 10.596% ; T: 89.404% | Imputed | 0.2 | 0.00003 | No |
| **rs34850343** | **X:6032382** | **A: 10.490% ; G: 89.510%** | **Genotyped** | **0.2** | **0.00003** | **Yes** |
| rs35005859 | X:6032730 | A: 80.530% ; C: 19.470% | Imputed | 0.17 | 0.000065 | No |

Cis eQTLs were precomputed in the GTEx database in a +/- 1Mb cis window around the transcription start site of the *NLGN4X* gene. This differed from the +/- 500kb window around the coding region of the *NLGN4X* gene analysed using the FUSION script. Interestingly, although the FUSION script took into account a smaller region than the GTEx analysis the eQTL SNPs identified covered a larger region (≈980kb in FUSION workflow compared to ≈280kb in GTEx workflow).

Three eQTL SNPS were found 3’ to the gene, within 55kb of the end of *NLGN4X*. The other 31 were found within introns of *NLGN4X* (Figure 2). This was in contrast to the eQTL SNPs identified by FUSION, which were mostly positioned 5’ and 3’ to the gene. No eQTL SNPs overlapped between the two inference processes.

### Quality Control of Identified eQTL Panels

The eQTL SNP-panels identified by FUSION and GTEx were each used to infer *NLGN4X* expression across two sample sets of twins, each one further divided into male and female datasets. This inference relies upon an overlapping genotype set between the identified eQTL-SNP panels and the twin datasets. Genotypes in the twin dataset were mainly imputed from a SNP array enabling a low missing rate and good variant coverage. However, each eQTL-SNP panel still had missing data in the test sample.

In the female cerebellum FUSION eQTL panel, genotype data were available in the twin datasets for 13 of the 15 eQTL variants (Table 1), all of which had an imputation quality metric (R-squared) above 0.7. The two missing eQTLs (rs12556975 and rs182391645) were able to be replaced with proxies selected by considering LD information in control populations. Both CEU and GBR control populations returned the same two SNPs; rs11094813 in place of rs12556975 (CEU r^2^ of 0.9 and GBR r^2^ of 1) and rs150758913 in place of rs182391645 (CEU r^2^ of 0.82 and GBR r^2^ of 0.85). These 2 proxy SNPs had been imputed in the twin dataset. However, inspection of the imputation information showed that one SNP (rs150758913) had a low quality metric (R-squared of 0.54). Therefore concordance was checked for all twin pairs for this SNP. All 43 monozygotic twin pairs assessed showed concordant genotypes for rs150758913 indicating that the genotype data were consistent across individuals. As this was the only proxy SNP available for rs182391645, it was added to the eQTL-panel.

All 15 SNPs in the female cerebellum FUSION eQTL panel were biallelic in the female twin dataset and Hardy Weinberg Equilibrium was maintained with p>10^-6^. As the SNPs had been imputed all genotyping rates were 100%. Further Quality Control checks revealed that this SNP-panel included variants that were in tight LD with each other and therefore may over-represent the effects of each single true functional eQTL variant. In particular, SNP pairs ‘rs12389068 and rs5961359’ and ‘rs17305593 and rs73182556’ were found to be in complete pairwise-LD with an r^2^ value of 1 in both twin female sample sets. To avoid over-representation, SNPs rs5961359 and rs73182556 were removed from further analysis leaving a final sample of 13 SNPs which were ultimately used to infer *NLGN4X* gene expression in the female twin dataset (Table 1).

All 34 of the variants included in the cerebellum GTEx eQTL-SNP panel were directly genotyped or imputed in the twin dataset (no proxies were required) and all were found to be biallelic in the twin dataset with Hardy Weinberg p-values >10^-6^ and genotype rates of 100%. Linkage disequilibrium analyses revealed that this eQTL-panel had a greater level of variant-relatedness than the FUSION cerebellum eQTL-panel with many SNPs showing r^2^>0.8. The panel was therefore pruned to generate an eQTL panel including 12 independent SNPs all with pairwise r^2^ <0.8 (Table 3).

### Inference of *NLGN4X* Expression in Independent Twin Dataset

Following the quality checks described above, the two final eQTL-SNP panels included 13 SNPs for the FUSION cerebellum panel and 12 SNPs for the GTEx cerebellum panel. Each of these panels were used to infer normalised *NLGN4X* expression levels in the female twin datasets. The GTEx panel was applied to females only to allow direct comparison between the inferred expression data. FUSION and GTEx cumulative expression values were plotted against frequency to compare the two methods of detection (Figure 3). The eQTL-SNP panel identified by the FUSION workflow gave a continuous scale with a more even distribution which was consistent between the two twin datasets (Figure 3a). Expression inferred from the GTEx eQTL-SNP panel appears to be more sporadic with most individuals having no change in expression levels and others randomly distributed above and below this level across a wide distribution (Figure 3b). Furthermore, the dataset used in the FUSION workflow could be separated into females and males for eQTL inference, making the results more reliable when analysing X chromosome SNPs. The distribution of the eQTL SNPs at the 5’ of *NLGN4X* in the FUSION workflow also appeared more convincing. This led us to conclude that the FUSION software gave the most reliable expression level output and would be the method to carry forward to the neurodevelopmental twin dataset phenotype analysis.

**Figure 3.**
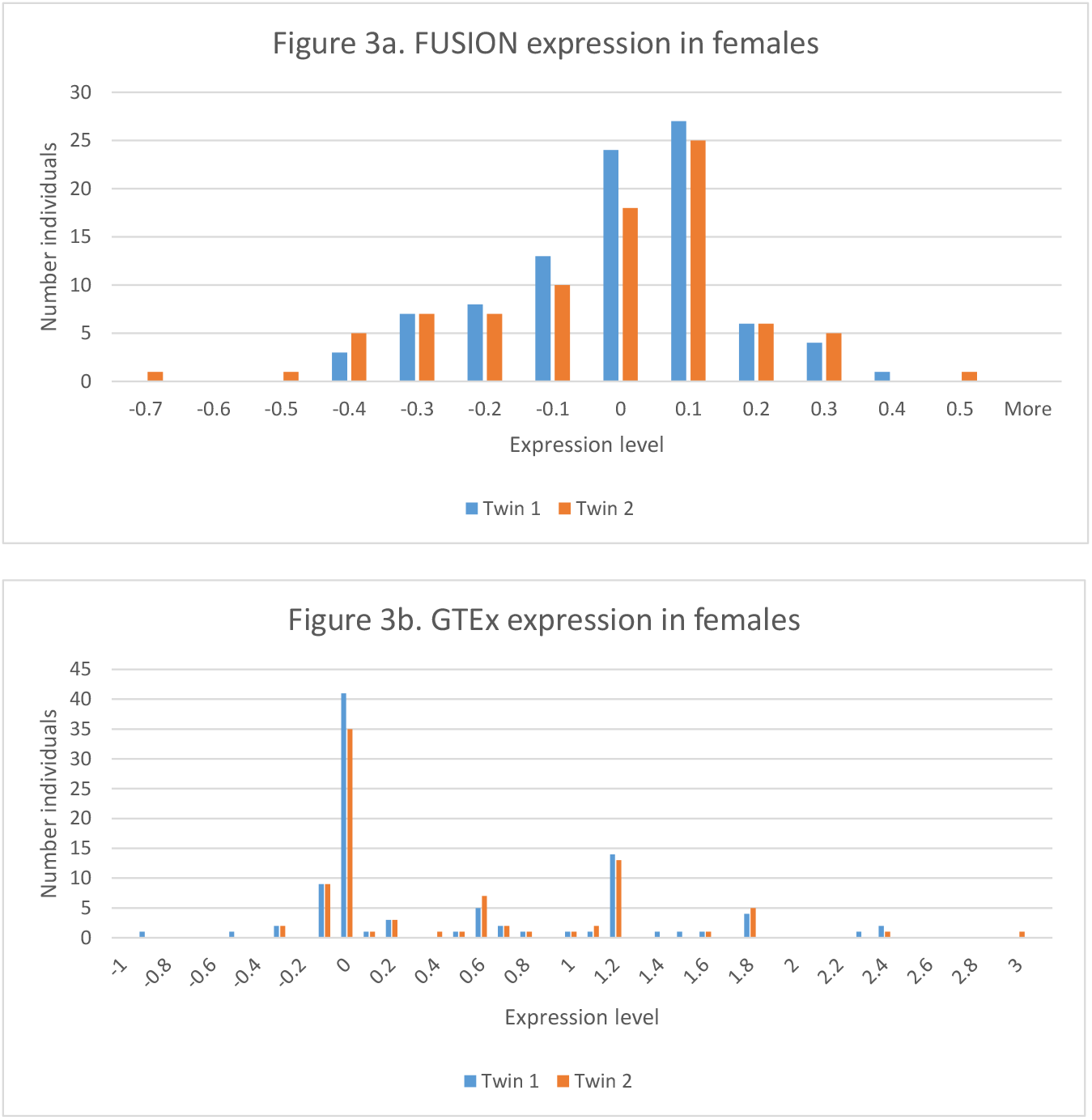
Comparison of the FUSION and GTEx cumulative expression outputs in the female Twin datasets.

### Association of Inferred *NLGN4X* Expression with Language Phenotypes

Inferred *NLGN4X* expression values from the FUSION cerebellar female eQTL-panel were investigated in a linear regression model for association to three quantitative measures of language skills and neurodevelopmental outcomes: language factor, nonword repetition and global neurodevelopment score. (Figures 4–6). Although trends of association were apparent in one twin set, in particular for global developmental score in twin set 1 (Figure 6), these were never replicated in the second twin sample (Figures 4–6).

**Figure 4.**
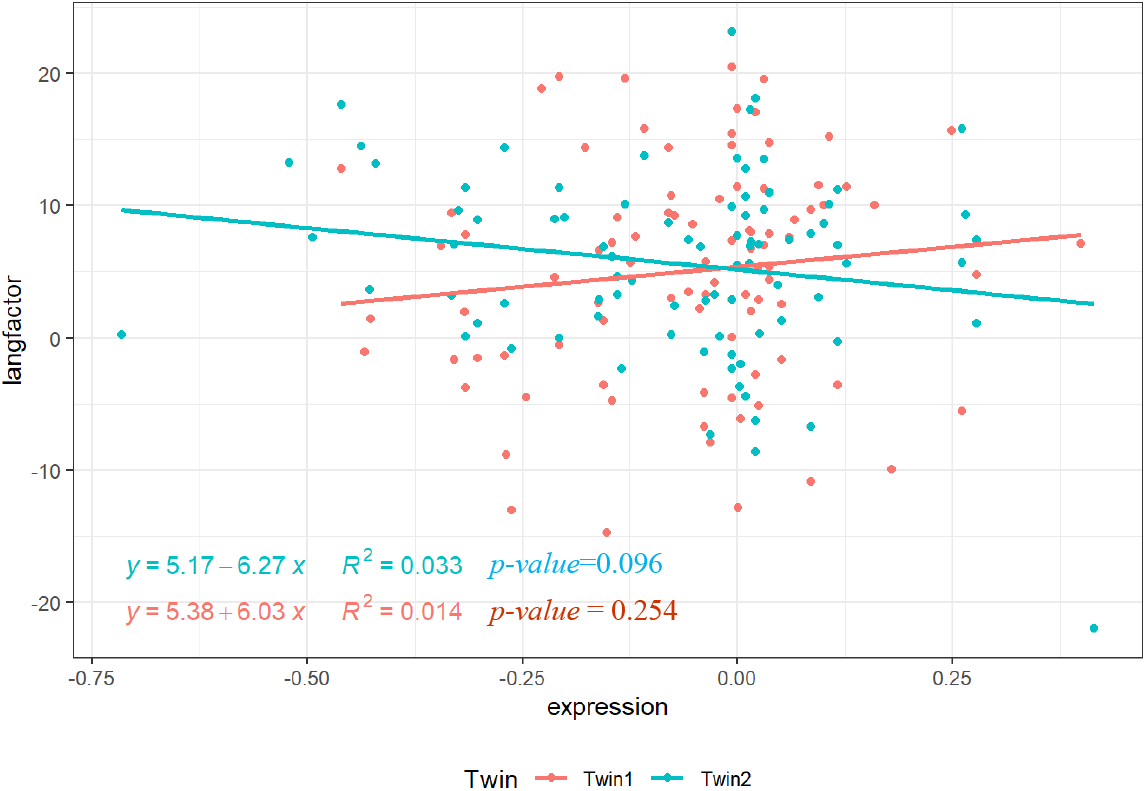
Twin 1 and 2 females - Language Factor vs *NLGN4X* expression in the cerebellum, from FUSION data.

**Figure 5.**
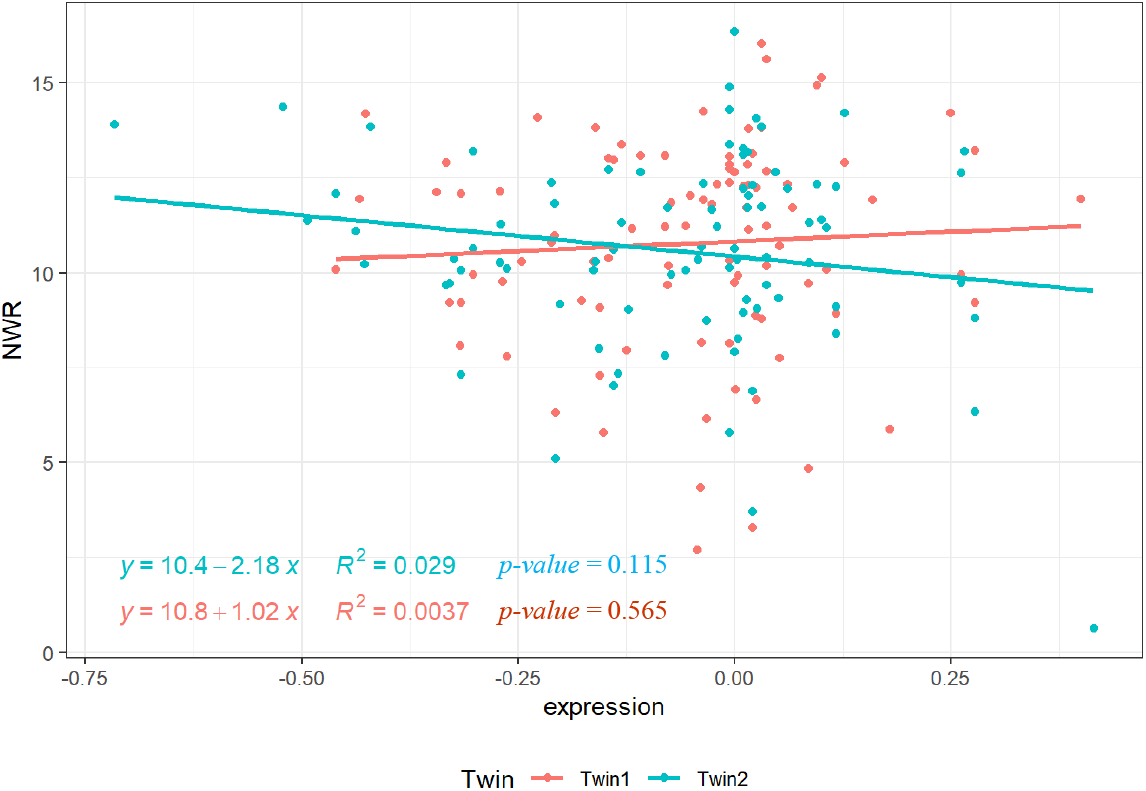
Twin 1 and 2 females - Nonword repetition vs *NLGN4X* expression in the cerebellum, from FUSION data.

**Figure 6.**
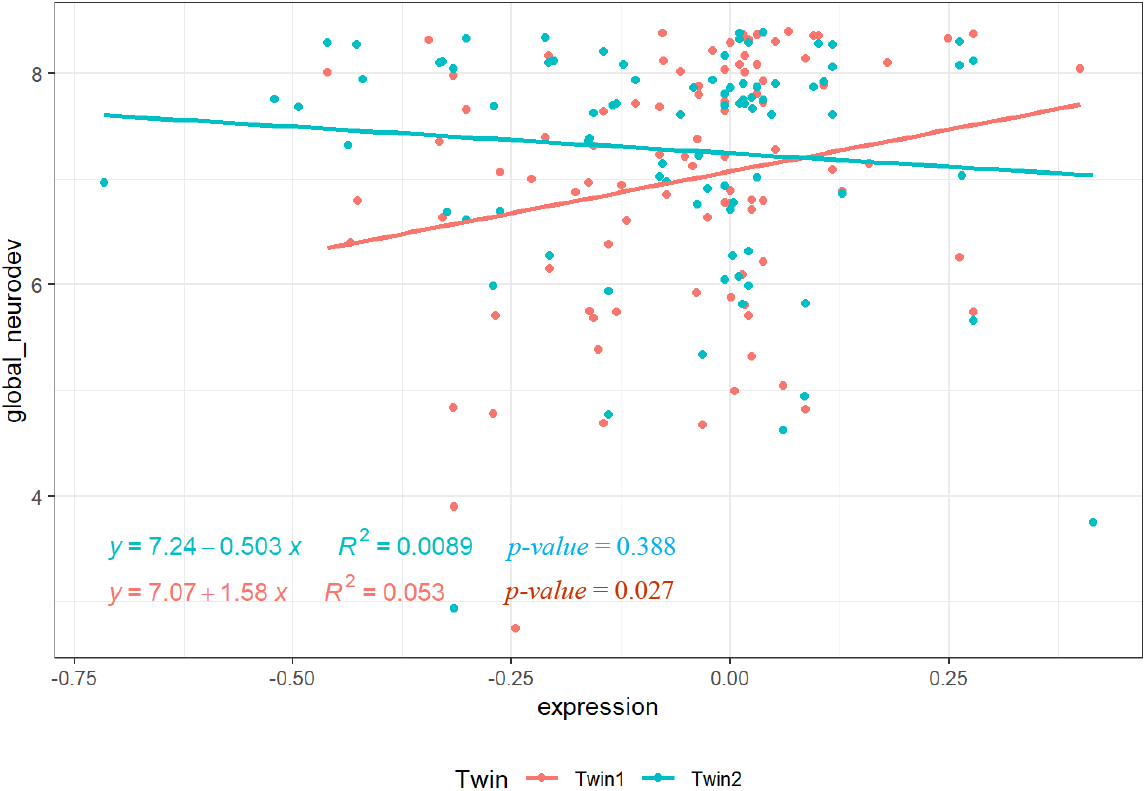
Twin 1 and 2 females - Global neurodevelopmental score vs *NLGN4X* expression in the cerebellum, from FUSION data.

## Discussion

In this study, we apply two methods of gene expression inference to two twin datasets. Recent research has seen an increase in publicly available genomic sequence data which capture common and rare genetic variation at the sequence level. However, these data are often not linked to gene expression, which may represent an important factor in susceptibility to complex disorders, because the investigation of gene expression requires different methods and sample types compared to the study of sequence data. Reliable inference of gene expression from genotype data therefore represents a potential tool in the interpretation of sequence variation. In this study, we focused upon a single candidate gene (*NLGN4X*) that has previously been associated with language development in relation to neurodevelopmental disorder. We hypothesised that altered expression of the *NLGN4X* gene may be associated with language ability and that this expression may be captured by the consideration of eQTL data.

We evaluated two methods of eQTL identification and capture – FUSION and GTEx. The FUSION workflow analysed genotype and expression files for correlations within individuals to identify putative eQTL SNPs. The GTEx workflow analysed a predetermined list of eQTL SNPs from the GTEx portal.

We identified several limitations to the application of these workflows. The FUSION workflow was initially difficult to run. We struggled to get a version of R that was compatible with all of the R packages needed to run the FUSION script. We ultimately used R version 3.4.1, which allowed a subset of models to be utilised to create a weighting. FUSION would not operate for X chromosome annotated SNPs therefore we analysed the female individuals alone so that the X chromosome SNPs could be analysed in the same way as autosomes. Genotypes had to be pruned to contain only SNPs present in the 1000 genomes LD reference data and were forced in Plink (Purcell et al., 2007) to have the same reference allele as the 1000 genomes. This was to make the genotype data compatible with the FUSION analysis. All of these steps were initially time consuming.

In contrast obtaining the eQTL SNPs from the GTEx portal was a simple download. However, there is no capability to distinguish male and female specific eQTLs on the GTEx portal, reducing confidence in results from the X chromosome. The FUSION method could be tailored to analyse females and males separately, which was particularly useful when analysing X chromosome SNPs.

The eQTL SNPs identified from the GTEx portal were very concentrated within the *NLGN4X* gene compared to the FUSION eQTL SNPs that were mainly 5’ and 3’. This was interesting as the GTEx *cis*-eQTL mapping window covered a larger region than that used in the FUSION workflow. Due to the concentration of SNPs there was high LD identified between the GTEx eQTL SNPs resulting in the 34 SNPs being filtered down to 12 independent SNPs. There was much less redundancy found within the FUSION eQTL SNPs, with only two being removed due to LD. For both methods the eQTLs in general did not fall in the 5’ region where promoters are expected to be. This shows the eQTLs identified may only be correlative with expression levels and not a direct mechanism of influence. No SNPs overlapped between the outputs from the two workflows indicating that the two methods for identifying eQTL SNPs differ substantially.

Overall, we found that the FUSION script allows a more fine grained imputation of gene expression, which allows for a better spread of expression values across eQTLs. The use of male data to impute X chromosome gene expression is limited due to the hemizygosity of males. The use of female data to impute X chromosome gene expression appears to be valid. Only a subset of brain tissues available can be expected to yield gene expression estimates and so investigations should be limited to two or three “most relevant” brain regions for the phenotypes. In this case the cerebellum was identified by the FUSION workflow as having eQTL SNPs for *NLGN4X*, this region of the brain has also been identified as having a role in language (Price, 2012; Silveri, 2021). Therefore, investigating correlations between expression levels of *NLGN4X* in the cerebellum and language phenotypes had an underlying biological theory.

No eQTLs were identified in the male neurodevelopmental twin set that had a MAF that allowed further analysis. No eQTLs were found using the FUSION software within a number of additional brain expressed genes and so correlations with language phenotypes could not be analysed.

In summary, this study did not identify any significant or consistent correlations between inferred *NLGN4X* gene expression and language phenotypes in a female neurodevelopmental twin sample set. The fact that no correlation was seen does not mean that the method does not work but may be a true negative. The obvious conclusion is that expression of *NLGN4X* in the cerebellum is unrelated to language development. Before discounting the relationship, however, we need also to note other possible reasons for null results. Our sample size was small and so would not be adequately powered to detect small effects. The analysis depends on the accuracy of imputed gene expression, which was inferred from a relatively small sample of deceased individuals. The sample size for inferring expression in the FUSION workflow differed between brain regions and was never more than 33 individuals for the females. We do not know how representative these individuals were of the general population. The age of the individuals at which the brain samples were taken will differ and the time of RNA extraction, which will reflect a single time point, and so the cells could have been performing differently at each time stage and producing different levels of RNA between individuals. The cause of death for the individuals could also have an effect on the levels of RNA. These analyses assume that there is a consistent relationship between genotype and gene expression, which may not always be the case. Previous studies have shown that expression of *NLGN4Y*, the *NLGN4X* homologue present on the Y chromosome, was associated with ASD phenotypes in XYY (Ross et al., 2015), whilst in this study we focused on XX females.

*NLGN4X* remains an interesting gene within neurodevelopmental and language phenotypes. Future work could use the FUSION software to look specifically at brain expressed genes within the *NLGN4* pathway.

## Acknowledgements

This work was funded by Wellcome Trust Programme Grants [082498], and European Research Council Advanced Grant [694189].

The Genotype-Tissue Expression (GTEx) Project was supported by the Common Fund of the Office of the Director of the National Institutes of Health, and by NCI, NHGRI, NHLBI, NIDA, NIMH, and NINDS. The data used for the GTEx workflow analyses described in this manuscript were obtained from: the GTEx Portal on 01/10/2019. The linked genotype and gene expression data used for the FUSION workflow were downloaded from dbGaP (Genotype-Tissue Expression (GTEx (https://gtexportal.org/home/)) (phs000424.v8.p2).

We offer warmest thanks to the families who took part in the study, and school staff who helped facilitate assessment arrangements. The study would not have been possible without the generous support of Prof Simon Fisher, Director of the Max Planck Institute for Psycholinguistics, who offered facilities for analysis of genetic material during the data collection phase of this study. We thank Katerina Kucera and Arianna Vino for their substantial work in processing DNA samples.

We thank the High-Throughput Genomics Group at the Wellcome Trust Centre for Human Genetics (funded by Wellcome Trust grant reference 090532/Z/09/Z) for the generation of the genotyping data.

Finally, the study would not have been possible without the hard work and dedication of eight research assistants who travelled around the UK to conduct assessments, including Nicola Gratton, Georgina Holt, Annie Brookman, Elaine Gray, Louise Atkins, Holly Thornton, Sarah Morris and Eleanor Paine.

## References

Bishop, D.V., and Scerif, G. (2011). Klinefelter syndrome as a window on the aetiology of language and communication impairments in children: the neuroligin-neurexin hypothesis. Acta Paediatr 100, 903–907.

Bishop, D.V.M., North, T., and Donlan, C. (1996). Nonword repetition as a behavioural marker for inherited language impairment: Evidence from a twin study. J Child Psychol Psyc 37, 391–403.

Danecek, P., Auton, A., Abecasis, G., Albers, C.A., Banks, E., DePristo, M.A., Handsaker, R.E., Lunter, G., Marth, G.T., Sherry, S.T., et al. (2011). The variant call format and VCFtools. Bioinformatics 27, 2156–2158.

Das, S., Forer, L., Schönherr, S., Sidore, C., Locke, A.E., Kwong, A., Vrieze, S.I., Chew, E.Y., Levy, S., McGue, M., et al. (2016). Next-generation genotype imputation service and methods. Nature Genetics 48, 1284–1287.

Gusev, A., Ko, A., Shi, H., Bhatia, G., Chung, W., Penninx, B.W.J.H., Jansen, R., de Geus, E.J.C., Boomsma, D.I., Wright, F.A., et al. (2016). Integrative approaches for large-scale transcriptome-wide association studies. Nature Genetics 48, 245–252.

Lonsdale, J., Thomas, J., Salvatore, M., Phillips, R., Lo, E., Shad, S., Hasz, R., Walters, G., Garcia, F., Young, N., et al. (2013). The Genotype-Tissue Expression (GTEx) project. Nature Genetics 45, 580–585.

Marien, P., Ackermann, H., Adamaszek, M., Barwood, C.H.S., Beaton, A., Desmond, J., De Witte, E., Fawcett, A.J., Hertrich, I., Kuper, M., et al. (2014). Consensus Paper: Language and the Cerebellum: an Ongoing Enigma. Cerebellum 13, 386–410.

Newbury, D.F., Simpson, N.H., Thompson, P.A., and Bishop, D.V.M. (2020). Stage 2 Registered Report: Variation in neurodevelopmental outcomes in children with sex chromosome trisomies: testing the double hit hypothesis. Wellcome Open Research 3, 85–85.

Nica, A.C., and Dermitzakis, E.T. (2013). Expression quantitative trait loci: present and future. Philosophical Transactions of the Royal Society B: Biological Sciences 368, 20120362–20120362.

Pai, A.A., Pritchard, J.K., and Gilad, Y. (2015). The Genetic and Mechanistic Basis for Variation in Gene Regulation. PLoS Genetics.

Price, C.J. (2012). A review and synthesis of the first 20years of PET and fMRI studies of heard speech, spoken language and reading. NeuroImage 62, 816–847.

Purcell, S., Neale, B., Todd-Brown, K., Thomas, L., Ferreira, M.A.R., Bender, D., Maller, J., Sklar, P., de Bakker, P.I.W., Daly, M.J., et al. (2007). PLINK: A Tool Set for Whole-Genome Association and Population-Based Linkage Analyses. The American Journal of Human Genetics 81, 559–575.

R Core Team (2013). R: A language and environment for statistical computing. R Foundation for Statistical Computing, Vienna, Austria.

Ross, J.L., Tartaglia, N., Merry, D.E., Dalva, M., and Zinn, A.R. (2015). Behavioral phenotypes in males with XYY and possible role of increased NLGN4Y expression in autism features. Genes Brain Behav 14, 137–144.

Silveri, M.C. (2021). Contribution of the Cerebellum and the Basal Ganglia to Language Production: Speech, Word Fluency, and Sentence Construction-Evidence from Pathology. Cerebellum 20, 282–294.

Wilson, A.C., and Bishop, D.V.M. (2018). Resounding failure to replicate links between developmental language disorder and cerebral lateralisation. PeerJ.

